# Trophic Transfer of Perfluoroalkyl Acids in a Periphyton-Mayfly-Zebrafish Food Chain

**DOI:** 10.1101/2025.04.09.647996

**Authors:** Matthew R. Farrell, David B. Buchwalter, Rebecca A. Weed, Jeffrey R. Enders, Antonio J. Planchart

## Abstract

Per-and polyfluoroalkyl substances (PFAS) are ubiquitous contaminants in freshwater ecosystems. Many PFAS are incorporated into food webs, with potential effects on ecological and human health. However, PFAS incorporation into the base of aquatic food webs remains poorly understood. The goal of this study was to quantify the uptake and trophic transfer of both legacy and current use PFAS compounds using a simulated freshwater food chain in a lab setting. Natural periphytic biofilms were placed into trays containing equimolar binary aqueous PFAS mixtures at environmentally relevant concentrations for five days. Following the initial exposure period, newly hatched mayfly larvae were introduced into each tray to feed on periphyton for most of their larval development. The mature larvae were then fed to zebrafish. All water and biota samples contained detectable levels of the tested PFAS. All PFAS were more concentrated in periphyton than in water, and four of six PFAS were further concentrated in mayfly larvae relative to periphyton. PFDA was the most accumulative in all biota. PFAS concentrations in zebrafish were significantly correlated with those in larval mayflies. Assimilation efficiencies in zebrafish were high (>70%) for all compounds. Bioaccumulation of PFAS in periphyton and mayflies was positively correlated with log K_OW_ and number of carbons.

## INTRODUCTION

Per- and polyfluoroalkyl substances (PFAS) are a class of over 14,000 anthropogenic chemicals that are used in numerous industrial applications (Williams et al, 2017). The C-F bond is the strongest bond in organic chemistry, which imparts uniquely useful properties to PFAS (O’Hagan, 2008). The fluorinated alkyl chain is highly stable and nonreactive, while charged functional groups present at the terminus of many PFAS allow for interactivity. Their amphiphilic structure and stability make PFAS powerful surfactants even in extreme temperatures and pH (Glüge et al, 2020). PFAS are widely used as processing aids in the production of PTFE (Teflon) and other fluoropolymers. They are also used in aqueous film-forming foams (AFFFs) to combat flammable liquid fires (Buck et al, 2011). PFAS are effective repellants of oil, water, and soil, leading to their use in protective coatings. These can be found in many consumer goods such as nonstick cookware, food packaging, and stainproof clothing or furniture (Gaines, 2023). PFAS exposure has been connected to adverse health outcomes such as immunosuppression, lipid dysregulation, thyroid disease, and cancer (Timmerman et al, 2020; Geiger et al, 2014; Winquist and Steenland, 2014; Barry et al, 2013).

PFAS have become environmental contaminants of concern due to their widespread use and persistence. Emissions from fluorochemical plants are major primary sources of PFAS pollution (Steenland et al, 2009; Sun et al, 2016; Zhang et al, 2024). The frequent use of AFFFs at airports and military bases is another common source (McGarr et al, 2023). Secondary sources of PFAS pollution include landfill leachate, agricultural runoff, and wastewater treatment plant effluent (Ghisi et al, 2019; Sun et al, 2016; Coffin et al, 2023). The stability that improves the utility of PFAS for commercial uses also makes them highly resistant to environmental and metabolic degradation. They can be transported great distances through rivers, groundwater, oceanic currents, and the atmosphere, resulting in detectable PFAS levels far from any known sources (Ahrens et al, 2023; Lyu et al, 2022).

Aquatic ecosystems are the ultimate sink for many PFAS in the environment, even when not released directly into water. PFAS in soil slowly leach into surface and groundwater following rain (Høisæter, 2019). Insoluble PFAS may degrade into perfluoro alkyl acids (PFAAs), whose anions are more hydrophilic compared to their precursors (Brase et al, 2021; Buck et al, 2011). Particularly high levels of PFAS are frequently measured in freshwater systems near point sources of pollution, as well as at sites far downstream. Well water near the Washington Works plant in Parkersburg, WV, was found to contain levels of perfluorooctanoic acid (PFOA) up to 13,300 ng/L (Hoffman et al, 2011). Further sampling revealed elevated levels of PFOA over 500 km downstream of the plant (Herrick et al, 2017). In North Carolina, total PFAS levels downstream of a Chemours plant exceeded 150,000 ng/L and included a mixture of newly identified novel PFAS compounds (Sun et al, 2016). An analysis of surface waters near US Air Force bases found a median concentration of 840 ng/L for the most common PFAS congeners. The PFAS mixtures in AFFF-affected waters are particularly high in long-chain sulfonic compounds such as perfluorooctane sulfonic acid (PFOS) and its precursors (East et al, 2021).

Many PFAS are bioaccumulative in freshwater biota. PFAS tissue concentrations are consistently elevated in fish, invertebrates, plants, and periphytic biofilms (Burkhard, 2021). Long-chain PFAAs are particularly accumulative, with PFOS being the predominant PFAS congener in most aquatic organisms (Ren et al, 2023; Brase et al, 2023; Simmonet-Laprade et al, 2019a). In the case of aquatic plants, this trend is at odds with terrestrial plants, which accumulate short-chain PFAAs more readily (Ghisi et al, 2019). There is a great deal of variance in bioconcentration factor (BCF) and bioaccumulation factor (BAF) values between different studies (Burkhard, 2021). Aqueous PFAS levels and physical characteristics of the study site can significantly affect bioaccumulation dynamics, which likely contributes to this variance (Lewis et al, 2022; Brown et al, 2023). The potential biomagnification of PFAS at higher trophic levels is also an area of concern. There is some evidence for the biomagnification of long-chain PFAAs in freshwater ecosystems, although there is little consistency between studies (Simmonet-Laprade et al, 2019a; Simmonet-Laprade et al, 2019b; Chen et al, 2018). Differences in PFAS accumulation between taxa likely contribute to these inconsistencies when attempting to assess biomagnification in entire trophic levels. The trophic transfer of precursor PFAS which then degrade into shorter-chain PFAAs may also play a significant role (Penland et al, 2020).

The bulk of research has focused on PFOA, PFOS, and a handful of other long-chain legacy PFAS. However, there are numerous other PFAS structures being widely used today. Data on the ability of these compounds to bioaccumulate and transfer up trophic levels are sparse. This represents a critical gap in the understanding of how these chemicals may affect natural food webs and the risk for human exposure.

To this end, we simulated a natural food chain using periphyton, mayflies (*Neocloeon triangulifer*), and zebrafish (*Danio rerio*). Periphytic biofilms form the basis of many freshwater food webs (Ijzerman et al, 2023). Mayfly larvae and other aquatic macroinvertebrates are common primary consumers that also serve as food sources for fish. Zebrafish are an established animal model that fills the role of secondary consumer. Here we have used this model to assess the uptake and trophic transfer of six legacy and novel PFAS (PFOA, PFNA, PFDA, PFHxS, PFOS, and NBP2) congeners at environmentally relevant concentrations. We quantified PFAS in water, periphyton, mayflies, zebrafish, and estimated assimilation efficiencies (AEs) in zebrafish. We then evaluated the relationship between PFAS structural characteristics and the previous endpoints.

## METHODS

### Mayflies and Periphyton

Naturally-grown periphyton and WCC-2 clonal strain mayflies (*N. triangulifur)* were obtained from the Stroud Water Research Center (SWRC; Avondale, PA). The WCC-2 strain was originally isolated from White Clay Creek in Chester County, PA by the SWRC (Sweeney and Vannote, 1981).

### Zebrafish husbandry

A wild type *Danio rerio* line was derived from AB wild type zebrafish obtained from the Zebrafish International Resources Center (ZIRC). All zebrafish lived in a carbon-filtered aquarium system at 28±1°C on a 14:10 hour day/night cycle. Tanks were kept at a density of 10 fish per liter or less. Salinity and pH were maintained at 800±50 µS and 7.5±0.5, respectively, via the addition of Instant Ocean salts and Iwaki aquatics pH solutions. Prior to experimentation, adult zebrafish were moved into individual tanks for 24 hours to acclimate and were fasted during this time. The NC State University Institutional Animal Care and Use Committee (IACUC, #24-231) reviewed and approved all procedures and uses of zebrafish.

### Chemical preparation

All PFAS were purchased from Synquest Laboratories: Perfluoro-n-octanoic acid (PFOA) (CAS: 335-67-1, product #2121-3-18), Perfluorooctane sulfonic acid (PFOS) (CAS: 1763-23-1, product #8169-3-08), Tridecafluorohexane-1-sulfonic acid potassium salt (PFHxS potassium salt) (CAS: 3871-99-6, product #6164-3-X4), Perfluorodecanoic acid (PFDA) (CAS: 335-76-2, product #2121-3-24), Perfluorononanoic acid (PFNA) (CAS: 375-95-1, product #2121-3-20), Nafion byproduct 2 (NBP2) (CAS: 749836-20-2, product #6164-3-3J). Working solutions were prepared in polypropylene carboys by diluting PFAS stock solutions in 20L of artificial soft water (ASW) (58 mg/L NaHCO_3_, 3.5 mg/L KHCO_3_, 22 mg/L CaCl_2_, 18 mg/L CaSO_4_(2H_2_O), 34 mg/L MgSO_4_(7H_2_O)). Each working solution contained an equimolar binary PFAS mixture specified in Table 1. Relevant physiochemical properties of each compound for this study are shown in Table 2.

**Table 1.** Target concentrations and actual measured concentrations for PFAS working solutions. Water samples were collected from experimental trays prior to addition of periphyton.

|  | PFAS | Target (nM) | Target (ng/L) | Measured±95%<br>CI (ng/L) | Error |
| --- | --- | --- | --- | --- | --- |
| **Mixture 1** | PFDA | 2.5 | 1285.2 | 507.3±30.9 | -60.5% |
|  | PFHxS | 2.5 | 1000.3 | 739.5±63.5 | -26.1% |
| **Mixture 2** | PFOA | 10 | 4140.7 | 4484.1±17 | +8.3% |
|  | PFNA | 10 | 4640.8 | 2642.6±84 | -43.1% |
| **Mixture 3** | PFOS | 10 | 5001.3 | 1954.1±522 | -60.9% |
|  | NBP2 | 10 | 4641.3 | 1769.5±427 | -61.9% |

**Table 2.** Names and evaluated characteristics of PFAS in this study. Central chain length was counted as the number of carbon, oxygen, and sulfur atoms in the longest straight chain. Log K_OW_ values marked with * are modeled estimates from Torralba-Sanchez et al. 2023. Log K_OW_ values marked with ** are modeled estimates from Wang et al. 2022.

|  | Central Chain<br>Length | Number of<br>Carbons | Molecular<br>Weight | Log K<sub>OW</sub> | Functional<br>Group |
| --- | --- | --- | --- | --- | --- |
| **PFDA** | 11 | 10 | 514.08 | 3.01* | Carboxylic Acid |
| **PFHxS** | 8 | 6 | 400.11 | 0.65* | Sulfonic Acid |
| **PFOA** | 9 | 8 | 414.07 | 1.79* | Carboxylic Acid |
| **PFNA** | 10 | 9 | 464.08 | 2.61* | Carboxylic Acid |
| **PFOS** | 10 | 8 | 500.13 | 1.69* | Sulfonic Acid |
| **NBP2** | 10 | 7 | 464.13 | 0.52** | Sulfonic Acid |

### PFAS exposures

Six replicate polycarbonate trays were each filled with two liters of the previously specified working solution. After the first exposure (PFDA / PFHxS mixture), the target concentration for each PFAS in subsequent experiments was increased from 2.5nM to 10nM to ensure detectable levels at all stages of the experiment. Exposure solutions were static and were not refreshed for the duration of the experiment. Trays were aerated using an air tube and placed under a grow lamp set to a 14:10 day/ night cycle. Trays were kept at room temperature, 23 +/− 1 °C. Five plastic slides covered in unexposed periphyton were submerged in each tray. The periphyton was left in place for five days. Water and periphyton samples were collected from each tray on days 0, 1, 3, and 5 of the exposure.

After five days of periphyton exposure to PFAS, 100 newly hatched mayfly nymphs were added into each tray. Mayflies were allowed to feed on periphyton in the trays for approximately 21 days, at which time we pooled larvae and collected three to five samples of 25-30 mayflies, dependent on the number of surviving larvae. Water and periphyton samples were also collected from each tray. Remaining mayflies were collected and transferred to a bowl of clean ASW before being fed to zebrafish. 10-20 adult zebrafish (number dependent on available mayflies) per trial were placed into individual 1 L tanks for 24 hours to fast and acclimate. Following the 24-hour fasting period, each individual fish was offered mayflies *ad libitum* within its tank until it stopped eating. After the feeding, half of the zebrafish were sacrificed and collected immediately. The remaining half were fasted for 24 hours after the feeding to allow gut contents to be purged, to make estimates of assimilation efficiency, in accordance with established zebrafish gut passage times (Cassar et al, 2018). Six fish were not fed any mayflies and were collected as controls for background PFAS levels present in the zebrafish.

### Sample collection

5 mL water samples were collected from the trays via polypropylene syringe with a 0.45-micron nylon membrane filter tip to prevent accidental collection of periphyton or detritus. Syringes and filters were rinsed with exposure solution three times before sample collection to reduce the impact of PFAS adsorption to the plastic. Water samples were stored in round-bottom polypropylene tubes.

Biological samples (periphyton, mayflies, and zebrafish) were collected into round-bottom polypropylene tubes and dehydrated in a drying oven at 60 °C for 48 hours. Both wet mass (pre-drying) and dry mass were recorded for all samples. For the first experiment (PFDA and PFHxS mixture), all biological samples were heat-dried. Downstream analysis with heat-dried zebrafish tissue proved challenging; therefore, zebrafish samples from the latter two exposures were freeze-dried instead.

### Sample Preparation

Samples were homogenized using a GenoGrinder bead mill. To this milled sample, internal standard and 0.01N NaOH (in methanol) were added. This mixture was vortexed and then centrifuged. Supernatant was removed, diluted with water and extracted via solid phase extraction (SPE) on a Supelco extraction manifold. Waters weak anion exchange (WAX) cartridges were first precleaned with 4 mL of 0.5% ammonium hydroxide and then conditioned with 4 mL of methanol prior to equilibration with 4 mL of water. Samples were loaded at a rate of about 1 drop per second. Cartridges were washed with 4 mL of 25mM ammonium acetate and then dried. Samples were eluted with 4 mL of methanol and then 4 mL of 0.5% ammonium hydroxide in methanol. Sample eluates were dried and then reconstituted with 1 mL of 50:50 methanol water (adding the methanol portion first) (Weed et al, 2022).

### Quantitative LC-MS Method

The samples were analyzed using an Agilent (Santa Clara, CA, USA) 6495C triple quadrupole mass spectrometer complexed with an Agilent 1290 Infinity II liquid chromatograph using method parameters similar to a previous report (Enders et al, 2022). Briefly, a 100 μL aliquot of each sample was injected onto a Kinetex F5 (2.1 × 100 mm, 100 Å; Phenomenex, Torrance, CA, USA) analytical column at 45 °C for separation. Aqueous (solvent A: water with 5% ACN and 0.1% formic acid) and organic (solvent B: ACN with 5% water and 0.1% formic acid) solvents were run at 500 μL/min using the following gradient: 0 min: 1% B, 2 min 1% B, 13 min: 70% B, 13.01 min: 99% B, 17 min: 99% B, 17.01 min: 1% B, 20 min 1% B. Additionally, an InfinityLab Poroshell HPH-C18 delay column (3.0 × 50 mm, 4 μm; Agilent, Santa Clara, CA, USA) was installed in the flow path of the LC to delay any potential PFAS contamination in the LC solvent from interfering with the sample analysis. A dynamic MRM method (with polarity switching) was employed for quantitation.

### Quantitative Data Analysis and Quality Controls

Results were quantitatively processed in Skyline (Maclean et al, 2010). For all compounds and internal standards, two mass transitions were monitored. Ion ratios between the two transitions were required to remain within 30% of the average ratio established by the calibration curve that was run on that given day, in order to be considered a valid result. Retention time had to match within ca. 6 s but small incremental shifts across the batch due to shifting chromatographic conditions were allowable. Sudden and drastic deviations from expected retention time would trigger a pause and rerun of effected samples. Expected peak shape was established by the calibration curve run with the batch and unknown samples with PFAS hits were required to have matching peak shapes compared to this calibration curve. A non-extracted calibration curve was analyzed with the samples as previously published using the following concentrations: 1; 2; 5; 10; 50; 100; 500; 1000; 2500; 5000; and 10,000 ng/L. Multiple quality control samples were used throughout the study. The sample preparation controls consisted of a neat positive control (light calibration mix spiked into distilled water at 800 ng/L with an identical spike of IS compared to samples) used to monitor extraction efficiency, and a negative neat control (distilled water spiked with only the IS mix) to monitor if any contamination occurred during the SPE step. The instrument controls were a non-extracted positive control (800 ng/L) to track instrument sensitivity over time and a negative control (IS only) to track any potential carryover. Once the on-instrument values were obtained, those concentrations were back calculated to the original sample concentration.

### Statistical analysis

PFAS concentrations at different timepoints in water and periphyton were compared to evaluate change over time using one-way ANOVA and Dunnett’s T3 multiple comparisons test. Linear regression was used to evaluate relationships between PFAS concentrations at different trophic levels. BAFs for mayflies were calculated as C_Mayflies(ng/kg dry mass)_/C_Water (ng/L)_ using mean aqueous PFAS concentrations from samples collected while mayflies were present. BAFs for periphyton were calculated as C_Periphyton (ng/kg dry mass)_/C_Water (ng/L)_ using mean aqueous PFAS concentrations from sample collected before the addition of mayflies. Assimilation efficiencies were estimated as C_Zebrafish_ (24 hours post-feeding) / C_Zebrafish_ (immediately post-feeding); pre and post gut clearance PFAS levels were compared using a student’s t test. Correlations between structural properties and BAF values were determined using Pearson’s correlation coefficients. All statistical analyses were performed using Graphpad Prism 10.4.1. Outlier detection for linear regression was done using Prism’s ROUT method with a false discovery rate of 1%.

**Figure 1.**
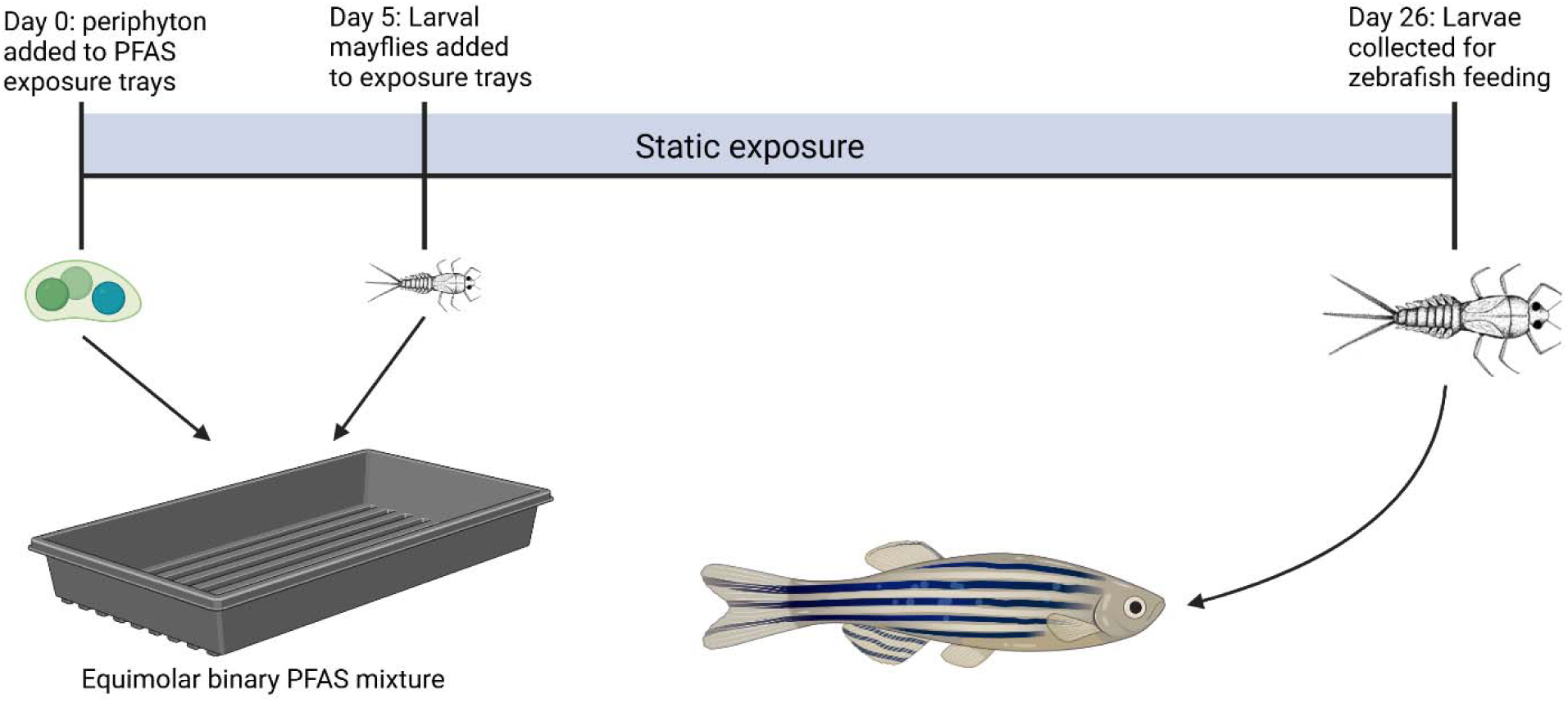
Visualized exposure protocol. Created in https://BioRender.com

## RESULTS

### PFAS uptake in periphyton and mayflies

Concentrations of each PFAS in water and periphyton during the experiment are shown in Figure 2. Initial PFAS concentrations in the periphyton were low (<10 ng/g dry mass) for all compounds. In all replicates, PFAS levels in periphyton increased significantly (p<0.05) by the 24-hour timepoint. In the same timeframe, aqueous PFAS concentrations decreased significantly for three congeners (PFDA, PFNA, PFOS) and non-significantly for the others. For both periphyton and aqueous PFAS levels, the largest changes were seen in the first 24 hours of the exposure. The majority of measured PFAS concentrations did not change significantly between 24 and 120 hours. PFOS increased in periphyton over that time period, while PFHxS decreased. Aqueous PFDA levels continued to decrease until the 120-hour timepoint. All PFAS were more concentrated in periphyton than in water, ranging from 52 ng/g (PFHxS) to 1362 ng/g (PFOS). PFDA was determined to be the most accumulative in periphyton (log BAF 3.46), followed by PFOS (log BAF 3.20) (Table 3). When comparing all tested PFAS, aqueous concentrations were not predictive of PFAS levels in periphyton.

**Figure 2.**
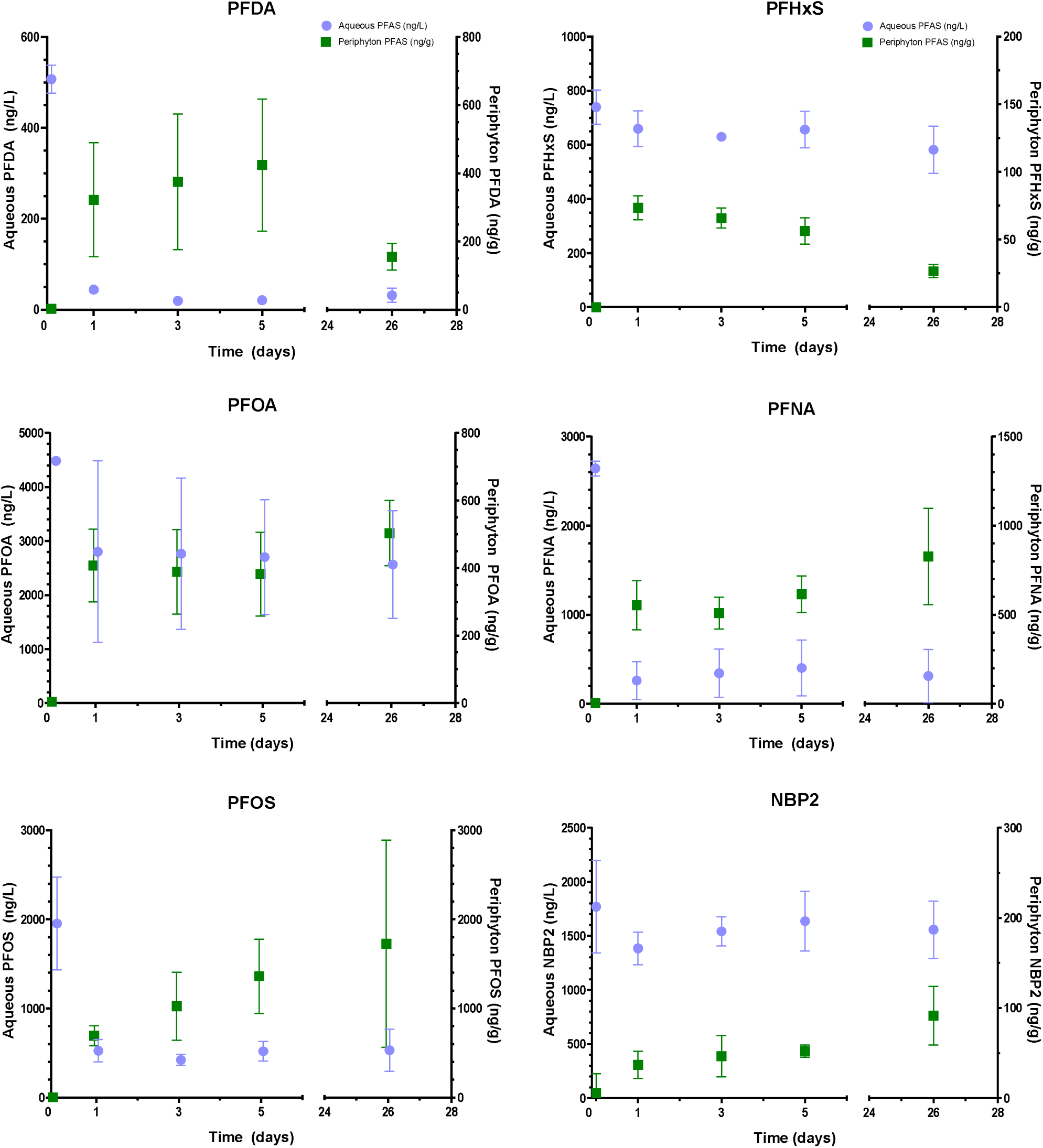
Concentrations of each PFAS in water and periphyton over 26 days. Bars represent 95% confidence intervals. Periphyton concentrations are ng/g dry mass. Aqueous PFAS were not refreshed during the exposures.

**Table 3.**
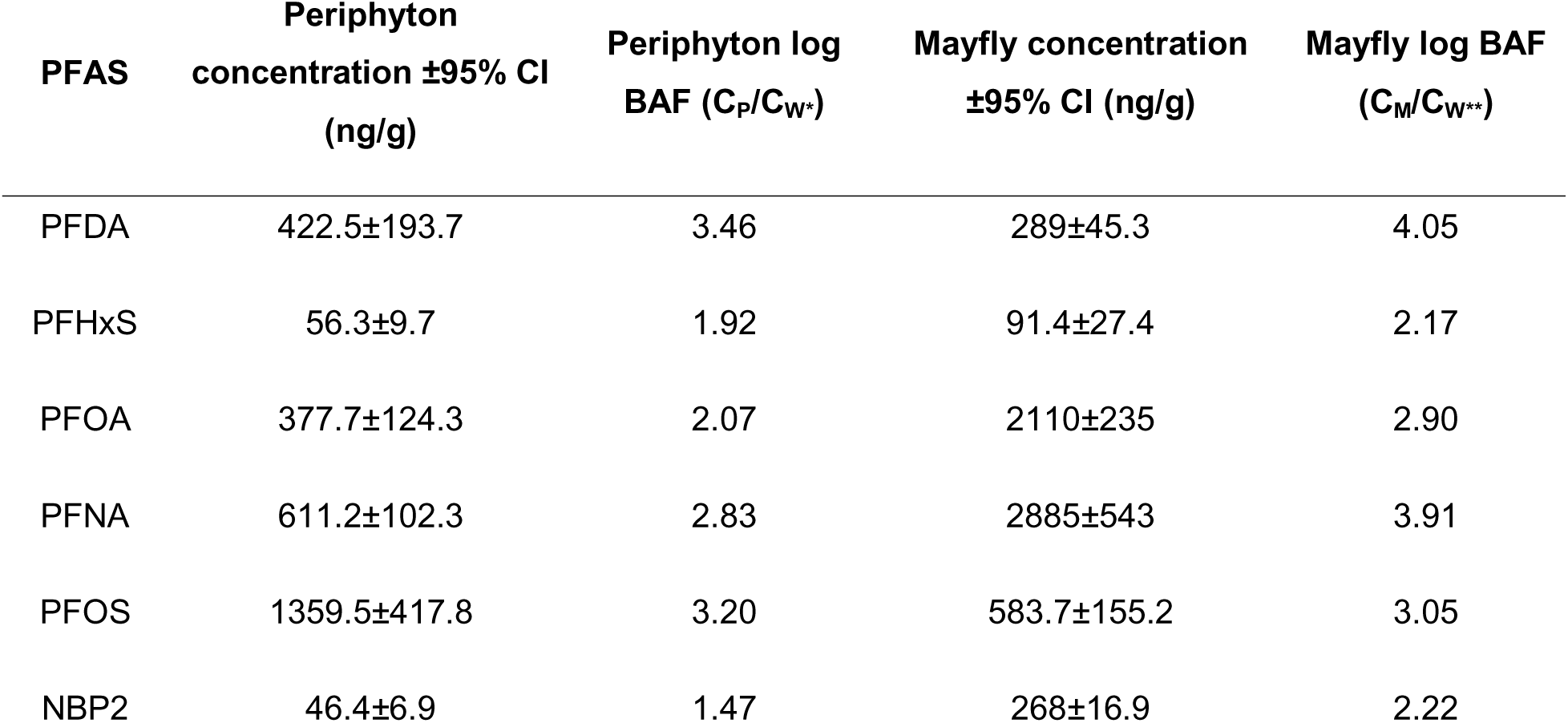
PFAS concentrations and bioaccumulation factors (BAFs) in periphyton and mayfly larvae. All biota concentrations were calculated using dry mass. Periphyton concentrations are means from day 5, prior to addition of mayflies. C_M_ and C_P_ are the concentration of the specified compound in mayflies and periphyton, respectively. C_W*_: Periphyton BAFs were calculated using mean aqueous PFAS levels from experimental days 0-5. C_W**_: Mayfly BAFs were calculated using mean aqueous PFAS levels from experimental days 5-26.

PFAS in water and periphyton were recorded again at 26 days, after mayfly larvae had grown to full size while feeding on periphyton in the trays. Relative to measurements taken at the 120-hour timepoint, before the addition of mayflies, there was no significant change in aqueous PFAS concentrations. PFDA and PFHxS had significantly reduced concentrations in periphyton at 26 days compared with the 120-hour timepoint. Nonsignificant increases were found for the other four PFAS (PFOA, PFNA, PFOS, NBP2), with PFOS and PFNA showing an upward trend.

PFAS concentrations in mayflies ranged from 91 ng/g (PFHxS) to 2885 ng/g (PFNA). PFDA was also the most bioaccumulative compound in mayflies (log BAF 4.05), followed closely by PFNA (log BAF 3.91) (Table 3). Aqueous PFAS levels did not significantly correlate with PFAS in mayflies. Four PFAS (PFHxS, PFOA, PFNA, NBP2) were enriched in mayfly larvae relative to periphyton, while two (PFDA and PFOS) were biodiluted. Periphyton PFAS concentrations were not predictive of concentrations in <u>mayflies</u>. However, PFOS was a major outlier when evaluating this relationship. With PFOS excluded, the remaining five congeners showed a strong positive relationship between periphyton and mayfly PFAS levels (Figure 3).

**Figure 3.**
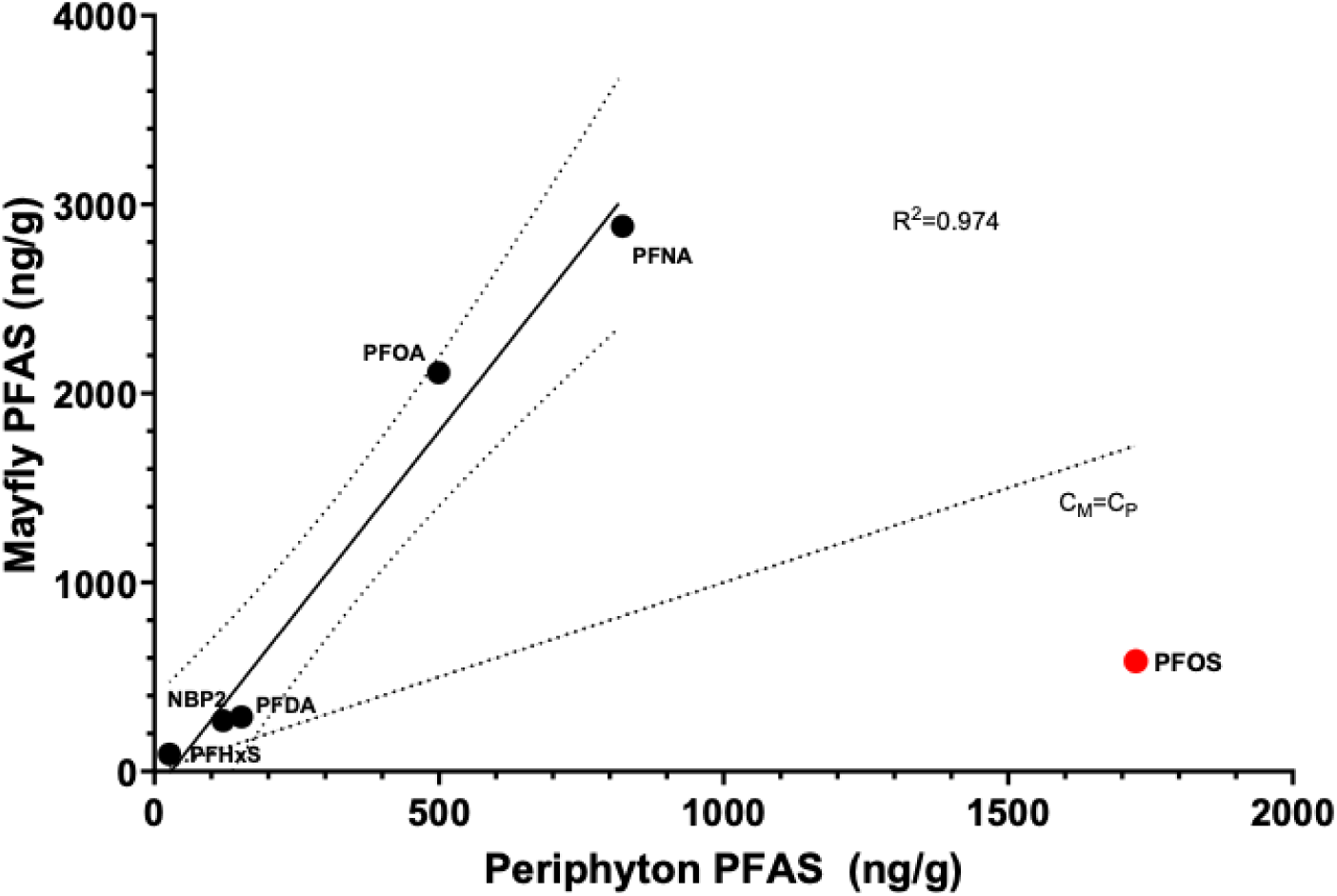
Linear regression trend and 95% CIs for PFAS in mayfly larvae vs PFAS in periphyton at experimental day 26. PFOS was determined to be an outlier (FDR=1%) and was excluded from linear regression. The dot-dashed line represents equivalent concentrations in mayflies and periphyton.

### PFAS uptake in zebrafish

Following a single feeding event, there were quantifiable PFAS in every exposed zebrafish. Mean measured concentrations in zebrafish ranged from 9.7 ng/g (PFHxS) to 91.7 ng/g (PFNA). Relative to PFAS levels in mayflies, PFDA was the most enriched in zebrafish (C_Z_/C_M_=0.137) and PFOA was the least (C_Z_/C_M_=0.025). PFAS body burden was significantly positively correlated with the number of mayflies ingested for five compounds (PFDA, PFHxS, PFNA, PFOS, NBP2) and showed a nonsignificant positive trend for PFOA (Figure 4). The number of mayflies ingested was not predictive of PFAS concentrations in zebrafish for any compound. However, total mayflies consumed correlated closely with body mass in zebrafish, counteracting trends related to dietary intake. Across the set of tested compounds there was a positive correlation between PFAS levels in mayflies and those in zebrafish (Figure 5). Male fish had significantly higher tissue PFHxS concentrations than females (14.3 ng/g vs 6.6 ng/g). This difference persisted when normalizing both groups for total dietary intake (0.08 ng PFHxS / mayfly ingested vs 0.04 ng/mayfly).

**Figure 4.**
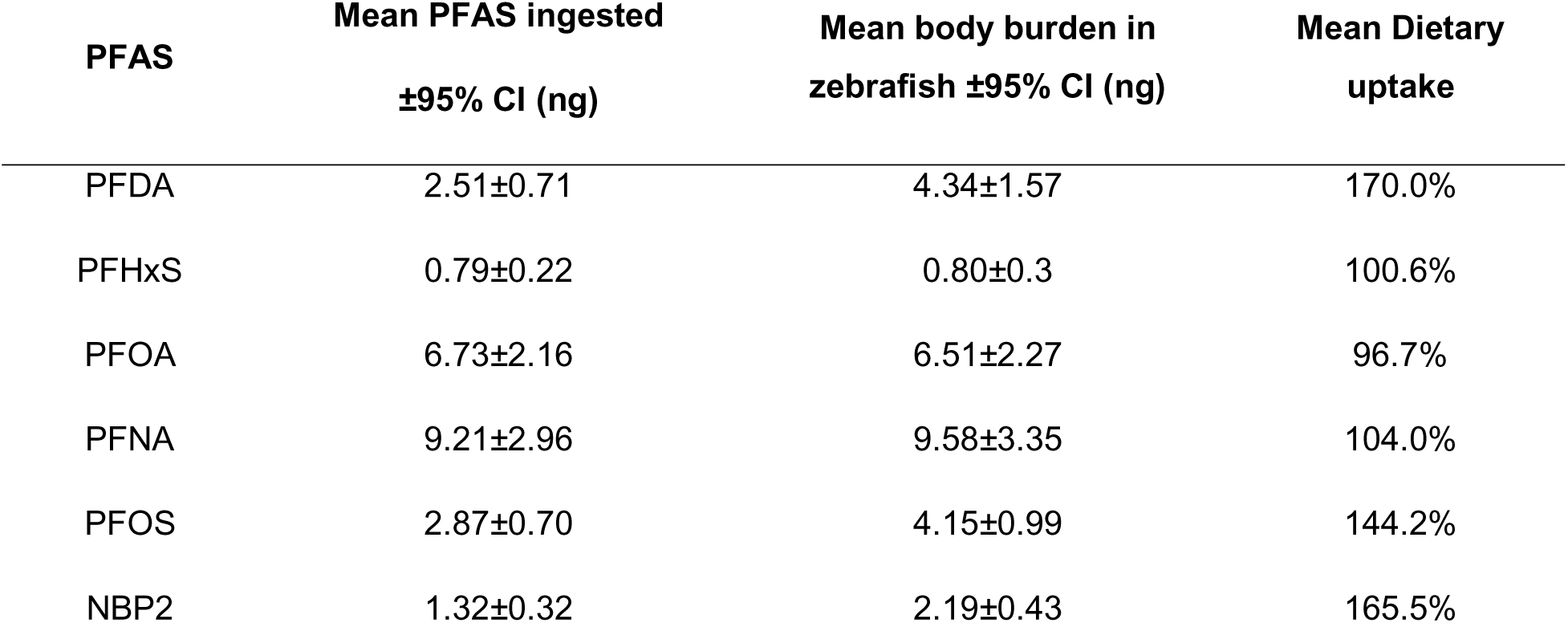
Linear regression and 95% CIs of total PFAS measured in zebrafish vs the number of mayflies ingested in a single feeding event. Red points were identified as outliers (FDR = 1%) and were excluded from linear regression. The slope of the linear regression line was significantly greater than zero for all PFAS except PFOA.

**Figure 5.**
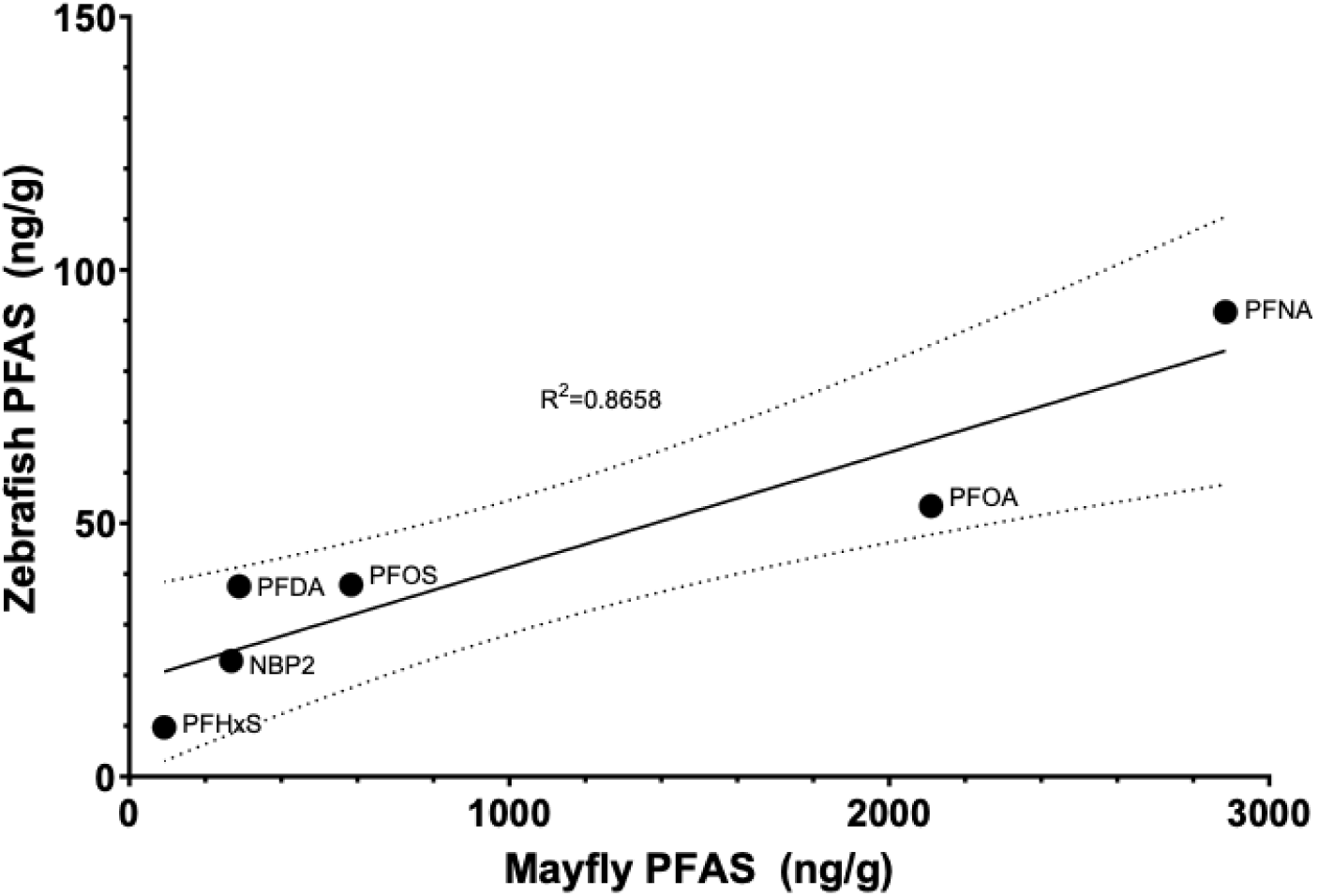
Comparison of PFAS concentrations in zebrafish and mayflies for all tested compounds following a single feeding event. All mass values are dry mass. The solid line was generated from linear regression. The dotted lines are 95% confidence intervals.

Estimated dietary intakes and body burdens for each PFAS are presented in Table 4. For the majority of compounds, the body burden in zebrafish exceeded the estimated intake from the mayfly feeding. The ratio of body burden to ingested PFAS varied greatly between individual fish (range: 0.47-3.59).

**Table 4.**
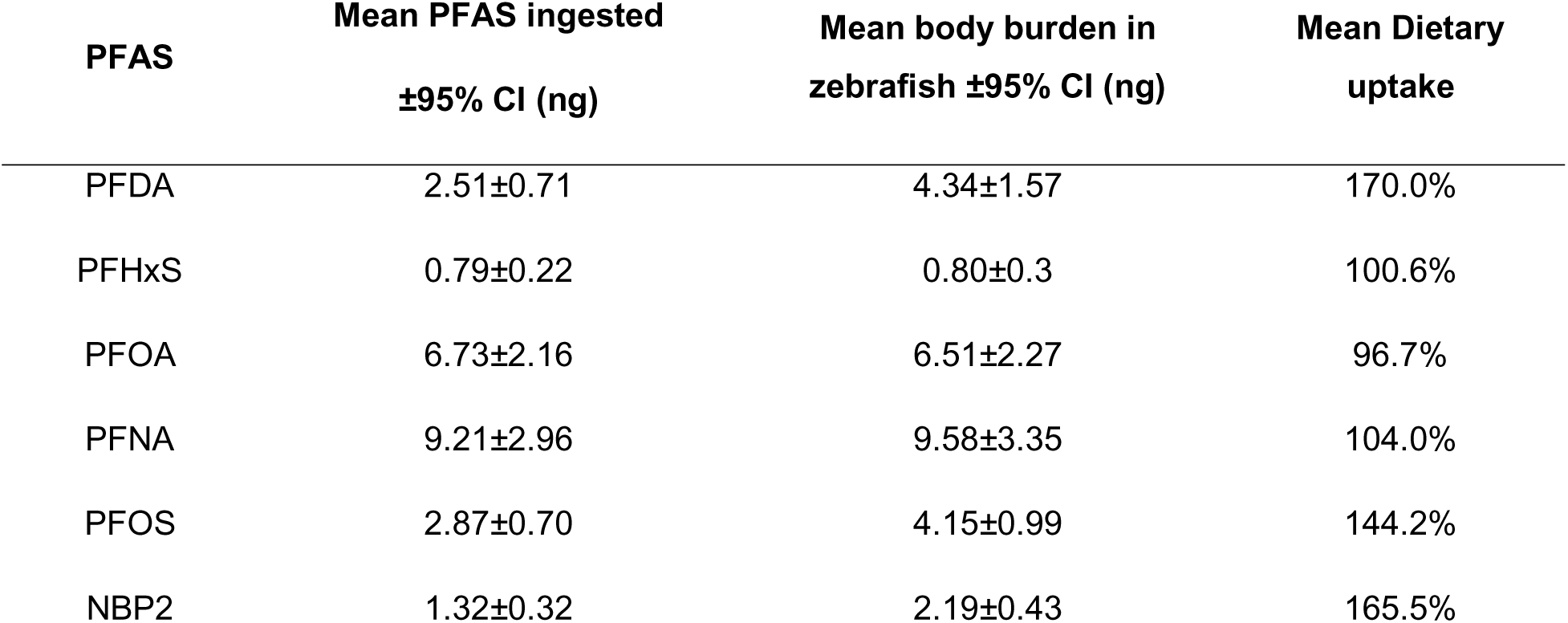
Total PFAS ingested per fish vs measured body burdens in zebrafish. Ingested PFAS was estimated by converting mayfly PFAS concentrations into PFAS per mayfly and multiplying by the number of mayflies ingested. Dietary uptake is presented as PFAS body burden / ingested PFAS.

| PFAS | Mean PFAS ingested<br>±95% CI (ng) | Mean body burden in<br>zebrafish ±95% CI (ng) | Mean Dietary<br>uptake |
| --- | --- | --- | --- |
| PFDA | 2.51±0.71 | 4.34±1.57 | 170.0% |
| PFHxS | 0.79±0.22 | 0.80±0.3 | 100.6% |
| PFOA | 6.73±2.16 | 6.51±2.27 | 96.7% |
| PFNA | 9.21±2.96 | 9.58±3.35 | 104.0% |
| PFOS | 2.87±0.70 | 4.15±0.99 | 144.2% |
| NBP2 | 1.32±0.32 | 2.19±0.43 | 165.5% |

Assimilation efficiency (AE) estimates for all PFAS are shown in Table 5. AEs were high for all compounds. Mean PFAS concentrations were not significantly different between fish that were collected immediately and those fasted for an additional 24 hours for any congeners. No sex-dependent differences in assimilation efficiency were found for any PFAS.

**Table 5.** Assimilation efficiency estimates for each PFAS. AE was derived by comparing PFAS concentrations (ng/g) in zebrafish collected immediately after dietary exposure and those fasted for 24 hours after exposure.

| PFAS | Post-feeding PFAS concentrations | PFAS concentrations after gut clearance | Assimilation Efficiency |
| --- | --- | --- | --- |
| PFDA | 43.18±16.15 | 32.01±12.26 | 0.74 |
| PFHxS | 10.00±5.26 | 9.43±5.55 | 0.94 |
| PFOA | 54.61±23.51 | 51.98±28.16 | 0.95 |
| PFNA | 80.38±32.15 | 105.9±73.19 | 1.32 |
| PFOS | 39.22±12.43 | 36.35±27.85 | 0.93 |
| NBP2 | 22.87±9.77 | 22.91±16.57 | 1.00 |

### Effects of PFAS structural properties on bioaccumulation

Log K_OW_ was the only characteristic significantly associated with bioaccumulation in both periphyton and mayflies (Figure 6). Number of carbons had a significant correlation with mayfly BAF and approached significance (p=0.059) in periphyton. Molar mass and central chain length were both positively associated with bioaccumulation, although neither was statistically significant. Bioconcentration and assimilation efficiency in zebrafish did not correlate with any assessed structural characteristics.

**Figure 6.**
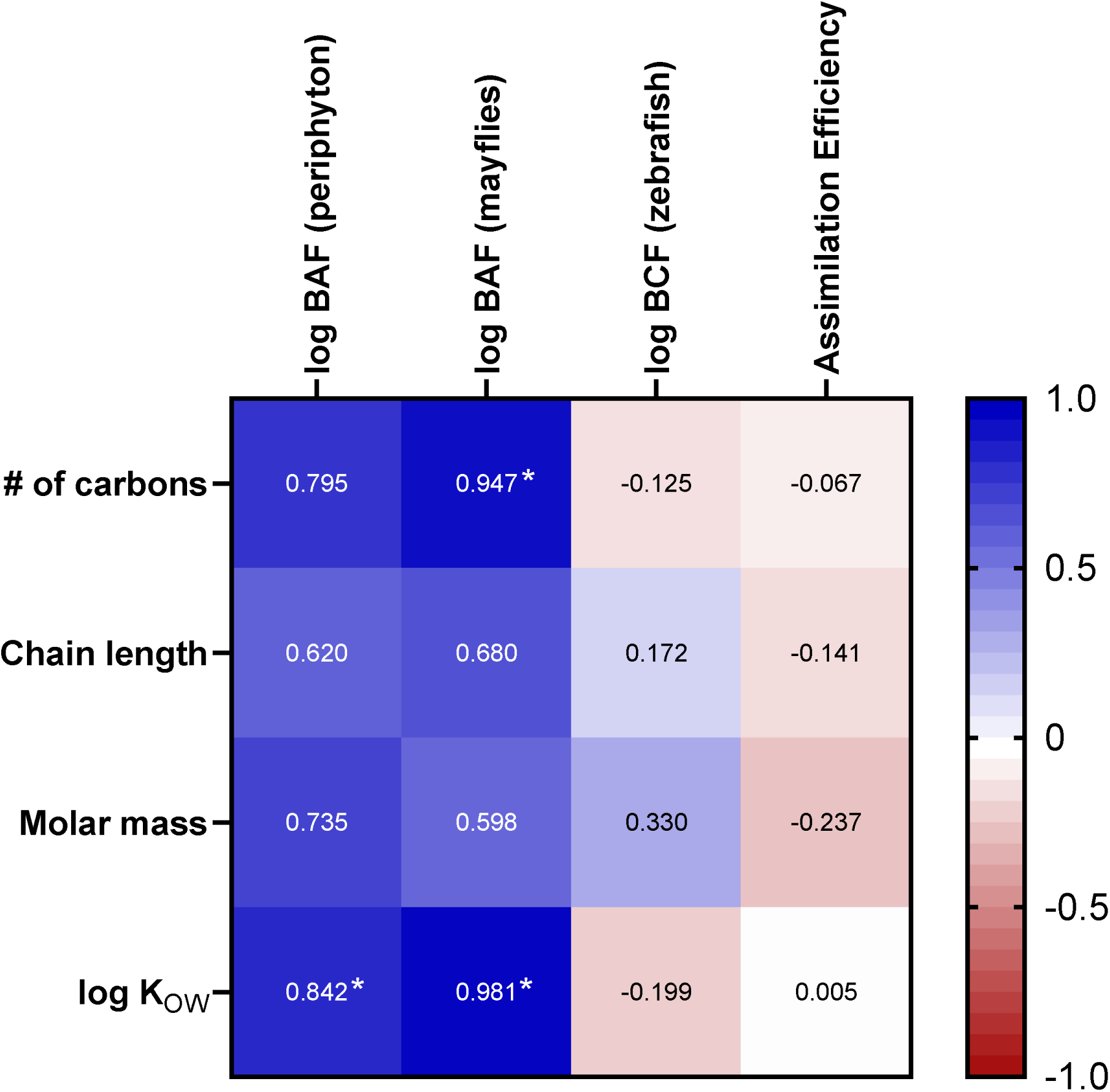
Pearson’s correlation coefficient for PFAS structural properties and bioaccumulation values. Statistically significant correlations (p<0.05) are marked with *.

## DISCUSSION

The sheer number of compounds and structural variety within the PFAS chemical class has made it difficult for the scientific community to adequately assess toxicity. Trophic transfer and bioaccumulative potential are important factors that contribute to human PFAS exposure through contaminated food sources. Our goal was to quantify the trophic transfer of PFAS in a multi-layered model food chain. This food chain captures multiple trophic levels that reflect real freshwater environments. Additionally, the species used require little space and grow quickly, allowing for the rapid testing of the trophic transfer for chemicals of interest. Similar models have previously been used to effectively model the trophic transfer of other environmental contaminants (Conley et al, 2014; Xie et al, 2010; Kim et al, 2012).

Our measured aqueous PFAS concentrations were lower than the target value for five compounds. Adsorption of PFAS to plastic labware is a commonly reported issue, and may have contributed to reduced aqueous PFAS levels. The impact of adsorption varies widely based on material, initial concentration, storage conditions, and the specific chemical. Studies examining PFAS adsorption to plastics used in this study (polypropylene, polycarbonate, nylon) have found that >50% of PFAS in water can be lost (Zenobio et al, 2022; Lath et al, 2019; Folorunsho et al, 2024). Efforts were made to minimize this effect during sample collection, but significant loss of aqueous PFAS likely still occurred.

All tested PFAS were detected in periphyton, mayflies, and zebrafish. All PFAS were more concentrated in periphyton than in the water, and four out of six PFAS were further concentrated in mayflies. PFAS concentrations in mayflies correlated more closely with those in periphyton than in the water, suggesting that dietary uptake was a significant contributor. Zebrafish were only ever exposed via a single feeding of contaminated mayfly larvae, indicating that all six PFAS were taken up via ingestion. Attempts to mass balance PFAS body burdens in zebrafish with estimated PFAS uptake from mayflies found that for PFAS levels in fish were significantly greater than their predicted total dietary intake for three compounds (PFDA, PFOS, NBP2). Dietary intake estimates were calculated based on the average mass of mayfly larvae and their PFAS concentrations, but there is considerable variation in the size of these larvae. If larval size affected zebrafish feeding behavior, our method may have underestimated dietary PFAS intake. It is also likely that in our study the efficacy of PFAS extraction prior to mass spectrometry varied widely between different matrices (Perovani et al, 2023). Mayfly larvae are less commonly analyzed than fish tissue, and lack standardized protocols. PFAS concentrations in mayflies may have been underestimated if extraction from insect tissue is less efficient.

There is very little literature on the bioaccumulation of PFAS in periphytic biofilms. These biofilms are diverse communities of organisms whose specific makeup can vary significantly between different freshwater environments (Levi et al, 2017). Additionally, it can be difficult to differentiate between internalization of PFAS and adsorption to the extracellular polymeric substances (EPS) that coat periphytic communities (Lewis et al, 2022). These factors make analysis of PFAS concentrations and accumulation in periphyton challenging. Several studies have calculated BAFs for periphyton in freshwater environments (Penland et al, 2020; Zhang et al, 2022; Munoz et al, 2018). The BAF values from this experiment were generally similar to those in the literature. Our results were in agreement with previous studies that have reported that long-chain PFAS and those with higher K_OW_ values are more bioaccumulative in biofilms (Zhang et al, 2022; Munoz et al, 2016). For legacy compounds of equal chain length (PFOA and PFOS), the sulfonic acid was more accumulative. To our knowledge, the bioaccumulation of NBP2 in periphyton has not previously been evaluated. NBP2 was the least accumulative PFAS in our test set, matching its low K_OW_ compared with similarly-sized PFAS compounds. Periphytic biofilms form the basis of freshwater food chains, thus they may be a significant contributor to the PFAS exposures of other biota. Our data on PFAS accumulation in biofilms contributes to filling an important gap in scientific knowledge.

We found several studies evaluating the bioaccumulation of PFAS in mayflies in natural environments. BAFs from this study matched closely with those presented in the literature. As in periphyton, log K_OW_ was found to be the feature most closely associated with increased accumulation, followed by number of carbons. These findings are consistent with other evaluations of PFAS bioaccumulation in mayflies (Ren et al, 2023; Brase et al, 2022). Aquatic insects or macroinvertebrates are commonly assessed as a single grouping in bioaccumulation studies. The variation in BAF values for grouped taxa are considerably larger than those focused on just mayflies (Penland et al, 2020; Simmonet-Laprade, 2019; Munoz et al, 2022; Yun et al, 2023; Koch et al, 2020; Lescord et al, 2015). Bioaccumulation can exhibit significant species-specific variance, and so community differences between sampled groups likely contributes to the wide range of BAFs (Groffen et al, 2023). For benthic macroinvertebrates, sediment PFAS levels may represent a more important exposure route than aqueous PFAS (Lewis et al, 2022). Additionally, dietary exposure can vary significantly between herbivorous and carnivorous invertebrates due to different PFAS accumulation dynamics in their food sources (Penland et al, 2020; Groffen et al, 2023). However, the general trends of bioaccumulation increasing with log K_OW_ and number of carbons are similar to those established in our experiment and in mayfly-specific studies.

Bioaccumulation in fish has been a greater focus of study compared with invertebrates and biofilms. As with other taxa, long-chain PFAS and sulfonic acids are particularly accumulative in fish (Lewis et al, 2022; Burkhard, 2021; Simmonet-Laprade et al, 2019; Lescord et al, 2015). This study did not find any correlation between PFAS uptake in zebrafish and any structural properties. However, in nature, fish would have both aqueous and dietary PFAS exposures where in our study, we focused exclusively on dietary uptake. We evaluated PFAS concentrations following only a single feeding, and so the departure from literature BAF trends is not unexpected. However, we did observe quantifiable trophic transfer of every tested PFAS in every fish, including the little-studied novel compound NBP2. The significant correlation between PFAS concentrations in mayfly larvae and in zebrafish lends further support to the relevance of diet as an exposure route for PFAS in freshwater environments. For all compounds, there was not a significant difference in PFAS levels between fish collected immediately after feeding and those given time to clear the gut. This suggests very high assimilation efficiencies for dietary PFAS in zebrafish. Previous research has also found the dietary assimilation of PFAS to be very efficient in fish (Martin et al, 2009; Xiong and Li, 2024; Kelly et al, 2009). Slow clearance of many PFAS congeners due to enterohepatic recirculation and partitioning to serum albumins are likely contributors to this high efficiency (Babut et al, 2017). Previous studies on gut transit time in zebrafish have been performed using larvae, so we cannot be certain that all gut contents were passed 24 hours after feeding. However, gut transit times in many other mature fish are less than 24 hours when measured in water temperatures comparable to those used in this study (Miegel et al, 2010; Navarro-Guillén et al, 2023).

PFAS exposure in freshwater animals is a function of both aqueous PFAS and dietary uptake. Many PFAS are bioconcentrated even in primary producers and low-level consumers, representing a significant source of PFAS uptake for consumers at higher trophic levels. Understanding the trophic transfer of PFAS is important to evaluate the risk these chemicals pose to freshwater ecosystems. It is also relevant for human exposure, as many fish from high trophic levels are important sources of protein in human diets. Ingestion is the main route of PFAS exposure for human populations, and seafood consumption has been positively correlated with serum PFAS levels (Hölzer et al, 2011; Christensen et al, 2017).

The environmental impacts of PFAS bioaccumulation in freshwater species are unclear. Numerous lab studies have established that PFAS are toxic to fish, invertebrates, and algae at environmentally relevant concentrations. PFAS exert a wide range of toxic effects including generation of reactive oxygen species, impaired photosynthesis, and developmental abnormalities (Gasparini et al, 2024; Liu et al, 2022; Ankley et al, 2021; Soucek et al, 2023). Invertebrates are generally more susceptible to PFAS toxicity compared with fish (Ankley et al, 2021). Less research has focused on the impacts of these chemicals on real-world freshwater ecosystems. A study of 30 streams in Belgium correlated PFAS body burdens with reduced biodiversity of macroinvertebrates (Byns et al, 2024)7 Fish samples from impacted environments have connected PFAS exposure to increased oxidative stress and biomarkers of altered immune function (Guillete et al, 2020; Piva et al, 2022). Further studies of natural communities are needed to elucidate which endpoints for PFAS toxicity are most relevant in freshwater ecosystems.

## CONCLUSIONS

This study demonstrated the trophic transfer of legacy and emerging PFAS through a three-level food chain. This model particularly sheds light on trophic transfer in periphyton and aquatic insects, important but understudied components of real-world food webs. Additionally, the short timeframe needed for these exposures permits rapid screening of trophic transfer potential in a controlled setting.

Trophic transfer in freshwater ecosystems is a significant contributor to dietary exposure risk for PFAS in humans. Trophic transfer and bioaccumulation are adequately understood for only a handful of PFAS in fish and not at all in aquatic insects and microbiota. An expansion of our studies to cover more diverse PFAS structures, concentrations, and mixtures would allow better understanding of the dynamics that drive trophic transfer in PFAS. This knowledge would aid in risk assessment of PFAS and the role of dietary exposure.

## ACKNOWLEDGMENTS

This work was performed in part by the Molecular Education, Technology and Research Innovation Center (METRIC) at NC State University. Funding for this project was provided by the National Institute of Environmental Health Sciences through award number P42ES031009.

